# Wing-pattern-specific effects of experience on mating behavior in *Heliconius melpomene* butterflies

**DOI:** 10.1101/2020.07.15.205435

**Authors:** Peyton A. Rather, Abigail E. Herzog, David A. Ernst, Erica L. Westerman

## Abstract

Many animals have the ability to learn, and some taxa have shown learned mate preference. This learning may be important for speciation in some species. The butterfly *Heliconius melpomene* is a model system for several areas of research, including hybridization, mate selection, and speciation, partially due to its widespread diversity of wing patterns. It remains unclear whether these butterflies can learn to prefer certain mates and if social experience shapes realized mating preferences. Here we test whether previous experience with a female influences male mate preference for two different *H. melpomene* subspecies, *H. m. malleti* and *H. m. rosina*. We conducted no-choice behavioral assays to determine if latency to court and whether males courted (vs no courtship) differed between naïve males and males with previous exposure to a young, sexually mature, virgin female. To test whether assortative courtship preference is learned in *H. melpomene*, males were either paired with a female who shared their phenotype or one who did not. Naïve *H. m. malletti* males courted assortatively, while naïve *H.m. rosina* males did not. Experienced *H. m. malleti* males reduced their courting relative to naïve males, suggesting that social experience with a sexually mature female that does not result in copulation may be perceived as a negative experience. In contrast, experienced *H. m. rosina* males exhibited similar courting rates to naïve *H. m. rosina* males. Our results suggest that social experience can influence male mating behavior in *H. melpomene* and that behavioral plasticity may differ across populations in this species.

## Introduction

Many of the behaviors and decisions that an animal makes are affected by its observations and capacity to learn. Learning can be defined as a set of processes that allows an animal to acquire, store, and use information gathered from the environment (Galef and Laland, 2005). Learning in animals is often complex and is likely the result of the social dynamics and settings of a species (Coussi-Korbel and Fragaszy, 1995). There is a substantial amount of evidence that animals have the ability to socially learn (Dukas, 1998). Some of the many behaviors that might be the result of social learning include food choices, predator avoidance, and mate preferences. For example, many species of fish have been observed to learn how to find food, how to recognize predators, and how to assess mate quality (Brown and Laland, 2003). This breadth of learning ability, however, is not limited to vertebrates (Dukas, 2008; Verzijden et al., 2012).

It is now understood that learning affects many essential activities of invertebrates, including predator avoidance and social interactions (Dukas, 2008, 2010). Particularly, many insects and spiders have shown the ability to learn mate preference. Studies on the wolf spider *Schizocosa uetzi* have shown that female social experience in their penultimate juvenile period can affect their mate choices as adults (Hebets, 2003). Female *Teleogryllus oceanicus* crickets modify their mate preferences after hearing attractive male songs (Bailey and Zuk, 2009), and female *Bicyclus anynana* butterflies learn preferences for enhanced male ornaments (Westerman et al., 2012). Male *B. anynana* also learn preferences for wing pattern elements in females (Westerman et al., 2014). Furthermore, work with *Drosophila melanogaster* fruit flies have shown that learning to be selective leads to a higher lifetime mating success than males who court indiscriminately (Dukas et al., 2006). Therefore, when it comes to mate preference and sexual behavior in insects it is often beneficial to learn.

Learning can potentially increase rates of assortative mating, which can lead to speciation through processes such as when young animals imprint on parents (Dukas, 2013). One such example of this is how cross‐fostering experiments in two subspecies of zebra finch demonstrated that assortative mating is due to imprinting. Birds in this study paired with mates that resembled their foster parents instead of their own phenotype (Irwin and Price, 1999). It has also been shown that mate preference can be learned in mature animals, such as male guppies and Syrian hamsters. These animals have demonstrated learning to discriminate against heterospecific mates after courtship interactions (Verzijden et al., 2012). This type of learning would help maintain speciation. With these studies in mind, we might expect that *Heliconius* butterflies, or other animals with high levels of speciation, might learn to court assortatively.

*Heliconius* butterflies have a long lifespan compared to other species of butterflies, which allows them to potentially mate multiple times (Gilbert, 1972). Therefore, the ability to learn in response to mating experiences could be advantageous. Studies have shown that male mate preferences evolve early in the speciation process in *Heliconius* within both intraspecific hybrid mating zones and conspecific polymorphic populations (Merrill et al., 2011a). These male mate preferences are based on wing color pattern cues, which are under natural selection to correspond to local mimetic environments (Gray and McKinnon, 2007; Kronforst et al., 2006). *Heliconius* is well known for its diversity in color patterns, and divergence in these color morphs is associated with speciation and adaptive radiation (*Heliconius*-Genome-Consortium* et al., 2012).

Here we take advantage of the social butterfly species *Heliconius melpomene,* whose widespread diversity of color patterns makes it an ideal model for studies on speciation and mating patterns (Jiggins et al., 2004). In this species, mimetic color patterns play a key role in species recognition, and mate preferences based on these patterns evolve alongside changes in wing pattern (Jiggins et al., 2004). Previous studies show that mimetic coloration in this species is important in choosing mates, and that these butterflies show assortative mating when choosing between their own and a different, closely related species (*Heliconius cydno*) (Jiggins et al., 2001). Furthermore, males often do discriminate between conspecific females with different wing patterns, and do not copy the mate preferences of conspecific males who have different wing patterns (Jiggins et al., 2004). However, it remains unclear whether individual *H. melpomene* males use past social experience with sexually receptive (or non-receptive) females to inform current mating decisions. The ability to learn mate preferences for intraspecies variation in wing pattern may be important for the initiation of assortative mating, reproductive isolation, and the speciation process.

Here we test whether experience impacts future male mate preference and courting behavior in two races of *H. melpomene* using three distinct *H. melpomene* color morph phenotypes (Figure 1). We had three alternative hypotheses: **1)** If males learn, then we predicted that experienced males would be more likely to court and have a shorter latency to court relative to naïve males. This type of learning is seen in *B. anynana*, where males exposed to dorsal hindwing spot number variation learn preferences for this trait (Westerman et al., 2014). **2)** If however male exposure to a female is somehow a negative experience, then we predicted that experienced males would court less often than naïve males. This type of learning is seen in *Drosophila* males. Studies have shown that naïve male *Drosophila* court virgin females persistently, while males previously exposed to an unreceptive female will then court virgin females much less vigorously (Siegel and Hall, 1979). **3)** If males are not able to learn, then courting was predicted to occur at random in both experienced and naïve males. An example of this is seen in the butterfly *Papilio polytes*, where a series of behavioral assays studying male preference for mimetic and non-mimetic females showed that there was no difference in initial and lifetime male preference, regardless of number of failed courtship attempts (Westerman et al., 2018).

**Figure 1.**
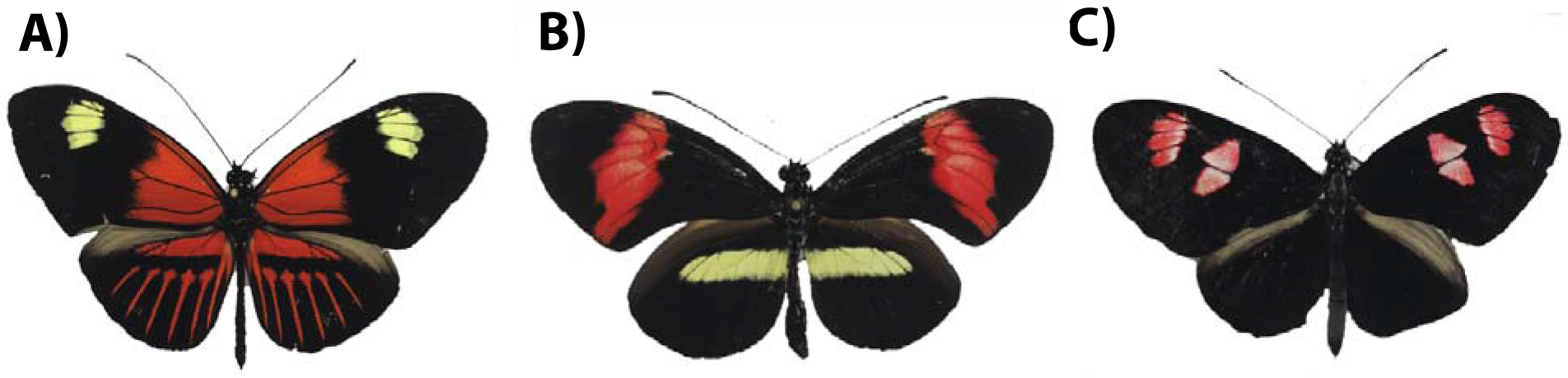
*Heliconius melpomene* phenotypes. A) *H. m. malletti*, B) *H. m. rosina,* and C) *H. m. plesseni*.

## Materials and Methods

### Study species and husbandry

*Heliconius melpomene* is a widespread neotropical butterfly found in Central and South America (Brower, 1994; Sheppard et al., 1985). The species is well known for its high diversity in color patterns, which play an important role in speciation (Jiggins et al., 2004). These color patterns, though diverse, are largely sexually monomorphic, with females and males of each morph having predominantly identical wing patterns. *H. melpomene* is often used as a model organism to study Müllerian mimicry and is co-mimetic with the species *Heliconius erato* (Jiggins et al., 2004). The many color patterns of *H. melpomene* have arisen through convergent evolution and selection for mimicry. The diversity of these patterns have allowed for the species to be used as models for studies on speciation and mating patterns (Jiggins, 2017). Here we take advantage of this species with different morphs that sometimes cohabitate in nature.

*H. melpomene* butterflies were either reared in a greenhouse at the University of Arkansas, or were obtained as pupae from Costa Rica Entomological Supply (Alajuela Apo. 2132-4050 Costa Rica.). Butterflies reared from the greenhouse colony came from a continuously breeding colony kept in two, phenotypic-specific, walk-in cages (143.51 × 110.25 × 219.71 cm). Caterpillars from the colony were given *Passiflora* plants *ad libitum*, and prior to pupation, plants containing caterpillars were removed from the breeding cages and moved to a separate 60.96 × 60.96 × 142.24 cm cage until butterfly emergence from pupa. Pupae obtained from Costa Rica Entomological Supply were reared on native *Passiflora* host plants in natural lighting as caterpillars prior to pupation and shipping. When at the University of Arkansas, they were hung in a 34.29 × 34.29 × 60.96 cm cage until emergence, and then maintained as adults in the Department of Biological Sciences Greenhouse at the University of Arkansas.

Butterflies were maintained at approximately 27°C, an average relative humidity of 71.5-85%, and a 13:11 light:dark cycle. The greenhouse was lit by Sun Blaze T5 high output 120-volt fluorescent light fixtures (containing UV wavelengths), in addition to natural sunlight, and the presence of UV light in the greenhouse was confirmed using an Ocean Optics Jaz spectrometer. After emergence, each butterfly was sexed, marked with a unique number on their hind wing (which does not harm the butterflies, see (Gall, 1984) for details), and then moved to 60.96 × 60.96 × 142.24 cm cages with food where they were kept until use in a behavioral watch. Males were placed into sex- and phenotype-specific cages, so they were isolated from both females and other wing patterns prior to behavioral assays. Females were placed into sex- but not phenotype-specific cages, so they were familiar with the wing patterns of males they were paired with in later behavioral assays. Each cage contained no more than 15 butterflies at any time and was visually isolated so that butterflies could not see individuals of the opposite sex (or phenotype in the case of males). Butterflies were fed BIRDS choice butterfly nectar (Birds Choice, Chilton, WI, USA), which is composed of glucose, fructose, calcium salt, halide salt, and amino acids. In addition, cages also contained *Passiflora* and *Lantana* plants for supplemental nectar and pollen. All behavioral watches were conducted in 60.96 × 60.96 × 142.24 cm BioQuip (Rancho Dominguez, CA, USA) observation cages, visually isolated from all other cages, between August 2017 and November 2019.

### Observational Experiment Time of Day Selection

To determine the time of day when the butterflies were the most active, we observed butterflies in colony cages for three consecutive days, between 6:00 am and 8:00 pm. Point counts were conducted every thirty minutes, where behaviors (*flight*, *walk*, *flutter*, *abdomen lift*, *bask* [defined by resting with wings held in open position], *rest* [defined by resting with wings held in closed position], *antennae wiggle*, *court*, and *copulate*) were recorded for each cage, followed by two ten-minute focal watches of one male and one female butterfly selected at random. Based on observations, we determined that butterflies were most active between the hours of 10:00 am to 2:00 pm.

### Behavioral Watches

All behavioral watches took place between 10:00 am and 2:00 pm, the time of peak *H. melpomene* activity in our greenhouse. Each watch consisted of a male aged ten or twelve days old, and a female between three and five days old. Watches were set up based on four separate treatments, N=15 per treatment per male phenotype. In this study, two male phenotypes were used (*H. m. malleti* and *H. m. rosina*) and three female phenotypes were used (*H. m. malleti*, *H. m. rosina*, and *H. m. plessini*) (Figure 1). To test whether males courted females with matching wing patterns faster than they courted conspecific females with dissimilar wing patterns, we tested latency to courtship and presence of courtship of naïve, 12-day-old *H.m. malleti* and *H.m. rosina* males matched with either females of their own phenotype or females of different phenotypes. To test whether previous exposure influences male latency to court, a 10-day-old (naïve) male was exposed to a female (with either a similar or dissimilar phenotype) until he either courted the female or 90 minutes passed without courtship. Afterward, the female was removed, and the male was returned to the all-male, phenotype-specific cage. On day 12, these males (i.e., experienced males) were exposed to a second female of the phenotype to which they had been previously exposed. The behavior of these males was then compared to that of naïve, 12-day-old males exposed to similar or dissimilar wing patterned females.

On the morning of a watch, a male was placed into an observation cage approximately two to three hours before the watch to acclimate to the new setting, and a female was added right before the start of a watch. Once a trial began, butterflies were observed for 90 minutes or until courtship took place. In the case that courtship did occur, time to court was recorded, and butterflies were not allowed to copulate. Behaviors were recorded to determine if any had an effect on mate preference. The number of incidents of each type of behavior (*flight*, *walk*, *flutter*, *abdomen lift*, *bask* (wings open), *rest* (wings closed), *antenna wiggle*, *sitting near*, and *court*) were recorded. Spectator Go BIOBSERVE (Fort Lee, NJ, U.S.A.) running on an Apple iPad was used to record time to court and all behaviors of the male and female during the testing period.

### Statistical Analyses

All statistical analyses were performed in JMP v. 14 (SAS Institute, Cary, NC, U.S.A.). We assessed whether latency to court was influenced by male experience or female wing pattern (similar or different from the male’s) using a GLM with male experience and female wing pattern as factors, as well as an interaction term. To assess if there was an effect of experience or female wing pattern on likelihood to court we ran a nominal logistic regression model using male experience and female wing pattern as factors, as well as an interaction term. Since we used two different morphs for our “different female” treatments (*H.m. plesseni* and *H.m. rosina* for *H. m. malleti* males; and *H. m. plesseni* and *H. m. malleti* for *H. m. rosina* males), we also tested whether there was an effect of female phenotype on male likelihood to court in our four different phenotype treatments using nominal logistic regression models. To test whether female behavior during a male’s first experience with a female had an effect on the observed courtship behavior in later interactions with females, we analyzed all behavioral data collected on day 10 watches (N=51 watches with behavioral data) and examined whether any of these behaviors were predictive of male courting on day 12. To do this we ran a principal components analysis on all the female behaviors and then ran logistic regression models on the first three principal components.

### Ethical Note

All *H. melpomene* butterflies were kept under laboratory conditions as defined by U.S. Department of Agriculture, Animal and Plant Health Inspection Service permit P526P-17-00343. Before being used in behavioral watches all butterflies were maintained in cages in a climate-controlled setting in conditions similar to those of their native habitat, and cages were inspected daily for ample food and appropriate conditions. All males and females used in a behavioral watch were either euthanized by freezing for use in future analyses or added to colony breeding cages where they were maintained in cages with ample food until natural death.

## Results

### Morph-specific effect of experience on likelihood to court

Experienced *H. m. malleti* males were less likely to court females than naïve *H. m. malleti* males, and were more likely to court *H. m. malleti* females than either *H. m. rosina* or *H. m. plessini* females (nominal logistic regression model, whole model χ^2^=14.935, p=0.0019, AICc=75.900, N=60; male experience χ^2^=8.85, p=0.009; female phenotype (same or different) χ^2^=6.85, p=0.009; interaction χ^2^=0.07, p=0.794) (Figure 2A). However neither experience nor female wing pattern influenced *H. m. rosina* male’s likelihood to court (nominal logistic regression model, whole model χ^2^=0.472, p=0.925, AICc=92.852; N=63, male experience χ^2^=0.313, p=0.576; female phenotype (same or different) χ^2^=0.126, p=0.722; interaction χ^2^=0.029, p=0.864) (Figure 2B). The effect of prior exposure to a female on likelihood of *H.m. malleti* to court two days later was independent of whether the male courted during the initial exposure period (nominal logistic regression model, whole model χ^2^=3.099, p=0.377, N=30; courted initially χ^2^=0.182, p=0.670; female phenotype χ^2^=2.325, p=0.127; interaction χ^2^=0.181, p=0.671). Male courtship response to females with different wing patterns was independent of the female wing pattern the male saw (*H. m. rosina* or *H. m. plesseni* for *H. m. malleti* males: nominal logistic regression model, whole model χ^2^=3.513, p=0.319, N=30; female phenotype χ^2^=0.683, p=0.409; male experience χ^2^=2.305, p=0.129; interaction χ^2^=0.391, p=0.531; *H. m. malleti* or *H. m. plesseni* for *H. m. rosina* males: nominal logistic regression model, whole model χ =0.188, p=0.979, N=32; female phenotype χ =0.002, p=0.966; male experience χ^2^ =0.387, p=0.844; interaction χ^2^=0.112, p=0.738).

**Figure 2.**
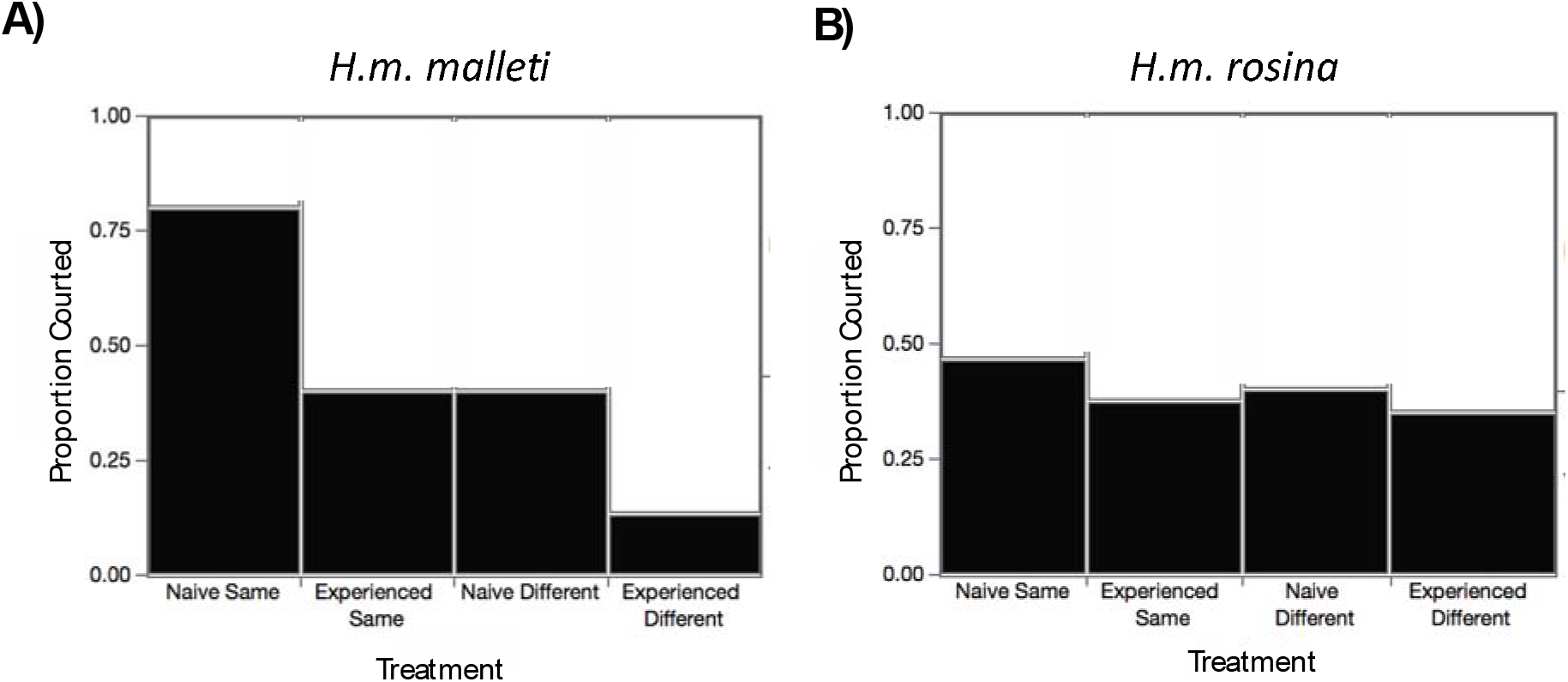
Lineage-specific effect of experience on male courtship. A) *H. m. malleti* reduce courtship after exposure to, but not copulation with, a female (N=60, male experience χ^2^=8.85, p=0.009; female phenotype (same or different) χ^2^=6.85, p=0.009; interaction χ^2^=0.07, p=0.794). B) *H. m. rosina* do not (N=63, male experience χ^2^=0.313, p=0.576; female phenotype (same or different) χ^2^=0.126, p=0.722; interaction χ^2^=0.029, p=0.864).

### No effect of experience or female wing pattern on latency to court

For those males that did court during the 90 minute observation period, there was no effect of experience or female wing pattern on male latency to court, for either *H.m. malleti* or *H.m. rosina* (GLM, *H.m.malleti*, whole model F ratio=0.391, p=0.761, N=23; male experience F ratio=0.375, p=0.548; female phenotype F ratio=0.599, p=0.449; interaction F ratio=0.038, p=0.848; *H.m. rosina*, whole model F ratio=1.663, p=0.205, N=25; male experience F ratio=0.200, p=0.659; female phenotype F ratio=3.22, p=0.087; interaction F ratio=1.452, p=0.242).

### Female Behavior Had No Effect on Future Likelihood to Court

We found no effect of female behavior during first exposure on male courtship rates during second exposure (Table 1 and 2).

**Table 1.** Loadings for principle components from PCA for female behavior during the training period for day 10 males.

|  | PC 1 | PC 2 | PC 3 |
| --- | --- | --- | --- |
| Flutter Count (Ct) | 0.513 | 0.091 | 0.046 |
| Fly Ct | 0.432 | -0.047 | -0.134 |
| Walk Ct | 0.441 | -0.171 | -0.010 |
| Lift Abdomen Ct | 0.061 | 0.216 | 0.817 |
| Bask Ct | 0.292 | 0.023 | 0.023 |
| Court Ct | -0.032 | 0.548 | -0.516 |
| Ant Wiggle Ct | 0.235 | -0.416 | -0.160 |
| Resting Ct | 0.19372 | 0.035 | 0.096 |
| Sitting Near Ct | 0.414 | 0.662 | 0.013 |
| % Variance Explained | 34.712 | 14.446 | 11.563 |
| % Total Variance Explained | 34.712 | 49.158 | 60.721 |

**Table 2.** Female behavior during early exposure did not influence male likelihood of courting in later female encounters. Test statistics and p-values from logistic regression models using composite behavioral variables PC1, PC2, PC3. N=51.

|  | **PC 1** | **PC 2** | **PC 3** |
| --- | --- | --- | --- |
| $\chi^2$ | 0.046 | 0.013 | 1.460 |
| **P-value** | 0.830 | 0.907 | 0.226 |

## Discussion

Our results show that male *H. melpomene* butterflies change their mating behavior in response to a social experience. This change in behavior is lineage specific, with *H. m. malleti* males, but not *H. m. rosina* males, exhibiting a reduction in likelihood to court after a social experience where they interact, but do not get to copulate, with a conspecific female. This effect was independent of the female’s wing pattern, though it did co-occur with a lineage-specific preference for assortative mating. *H. m. malleti* males courted *H. m. malleti* females more often than *H. m. rosina* and *H. m. plesseni* females in no-choice assays, while *H. m. rosina* males courted all female *H. melpomene* wing patterns equally often. Male likelihood to court was not significantly influenced by female behavior, and when males did court, experienced males did not court faster than naïve males.

*Heliconius* butterflies have many of the characteristics often found in species where past experience informs future social behavior; thus, our finding that *H. melpomene* males modify their mating behavior in response to experience, while novel, may not be unexpected. *H. melpomene* butterflies are relatively long-lived (up to 6 months in nature) (Gilbert, 1972), highly social (they roost in groups at night) (Mallet and Gilbert, 1995), and learn food sources and color cues (Toure et al., 2020). They have large brains (Montgomery et al., 2016) and are both physically larger, and longer lived than the butterfly *Bicyclus anynana*, which also uses past experience to inform current mating behavior (Dion et al., 2020; Westerman et al., 2012; Westerman et al., 2014). However, the negative effect of the pre-mating social exposure, and the wing-pattern-specific response to this pre-mating social exposure, were unexpected.

We initially hypothesized that early exposure to a female would prime the males to court faster and more often upon second exposure to females. This was based on previous findings in *B. anynana*, where naïve males do not exhibit a mate preference, but males with previous social experience do (Westerman et al., 2014). However, we found instead that early exposure to a female was a negative experience for *H. m. malleti* males, as they tended not to court on the repeat trial when exposed to a female of the same wing pattern and age as they had previously seen. One possible cause for this negative response could be that males who do court on day 10 are pulled from the watch upon courtship and are not allowed to copulate. This type of negative learning occurs in *Drosophila melanogaster* males, where previously unsuccessful males are reluctant to court females who smell similarly to females who previously rejected them (Griffith and Ejima, 2009; Siegel and Hall, 1979). *Heliconius* butterflies do have species-specific olfactory signals which are used in mating decisions (González-Rojas et al., 2020; Mérot et al., 2015), and olfactory signals have been shown to influence visual learning in *B. anynana* butterflies (Westerman and Monteiro, 2013). It would be interesting to see if olfactory cues play a similar role in *Heliconius*.

Although males did not learn to prefer certain phenotypes, avoidance learning from a negative experience could be beneficial to these males. *D. melanogaster* males have demonstrated learning to reduce courting females of the species *Drosophila simulans*, as these females typically reject mating attempts by male *D. melanogaster* (Dukas, 2004). Heterospecific courting can be costly for males because they could be wasting time and energy courting females that are likely to reject them (Dukas, 2009). This suggests that learning to avoid unreceptive females, or learning mate preference in general, may be beneficial relative to indiscriminately courting females. Negative learning in *H. m. malleti* may therefore lead to higher lifetime mating success, particularly if females who reject them once are likely to reject them again, and re-encounter rates are high due to nighttime social roosting.

It is worth noting that a characteristic of the social experience we used, the removal of a male as soon as he initiated courtship, resembles a component of the experimental design historically used in *Heliconius* butterfly mate choice trials. In many studies, males are allowed to approach a female and initiate courtship, and then they are physically removed from the female before given an opportunity to copulate (Chamberlain et al., 2009; Kronforst et al., 2006; Merrill et al., 2019; Merrill et al., 2011b). These males are then tested repeatedly, and past experience is often not accounted for when male preference is assessed, assuming that past experience does not inform present courting decisions. This assumption is partially based on a previous study showing that exposure to conspecific females with different wing patterns does not induce a preference for those wing patterns (Jiggins et al., 2004). However, this earlier study did not test for a negative effect of exposure on male preference or courtship behavior. Our results support the previous finding that prior exposure does not induce a positive preference, but suggest it instead may be a negative experience, at least for *H. m. malleti*. This wing-pattern specific response to experience should be seen as a cautionary tale for future comparative research with *Heliconius* butterflies, as repeated trials may be experienced differently by different lineages, which could confound interpretation of results. It also highlights the importance of checking for both positive and negative valence when testing the presence of learning.

The presence of both a lineage-specific response to prior experience and a lineage-specific presence of assortative courtship suggest that *H. m. malleti* and *H. m. rosina* may experience different mating-related selective pressures. Maintaining the capacity to learn can be energetically costly, and is often associated with fitness trade-offs, such as reduced fecundity (Kotrschal et al., 2013; Snell-Rood et al., 2011), reduced lifespan (Burger et al., 2008; Kotrschal et al., 2019), or extended development time (Kolss and Kawecki, 2008). Though conspecifics, *H. m. malleti* and *H. m. rosina* rarely co-occur in nature (Brower, 1996). The stronger innate preference and response to prior social experience exhibited by *H. m. malleti* suggest that *H. m. malleti* butterflies may, on average, co-occur with a more diverse *Heliconius* butterfly community than *H. m. rosina* butterflies, as is suggested by previously published range maps (Brower, 1996; Sheppard et al., 1985). This could lead to the maintenance of male assortative courtship and response to prior social experience in *H. m. malleti*, as limiting courtship efforts to those most likely to be successful would be energetically adaptive in an environment with many unreceptive females (Dukas et al., 2006). Assortative courtship and response to prior social experience may not be as strong in *H. m. rosina* as a result of either differences in generational exposure to polymorphic conspecifics, or differences in female receptivity. If most *H. melpomene* females are receptive to *H. m. rosina* males, independent of female wing pattern, there would be little pressure for males to maintain an assortative preference among conspecific wing patterns, or to maintain the ability to use past social experience to inform current courting behavior.

*H. m. malleti*’s response to prior experience could be associated with its assortative preference or a high social learning capacity. If *H. m. malleti* males have a stronger innate assortative preference than *H. m. rosina* males, it is possible that the initial exposure to a female creates a stronger negative memory for *H. m. malleti* males than for *H. m. rosina* males. Aversive signals are easier than appetitive signals for *D. melanogaster* to learn, and this is hypothesized to be due to the type of response to the initial cue (Schwaerzel et al., 2003). Alternatively, *H. m. malleti* might have a higher learning capacity (social or otherwise) than *H. m. rosina*. *Heliconius cydno* and *Heliconius melpomene* have different sized brain neuropils associated with sensory processing (Montgomery et al., 2020); it would be interesting to see if similar neuropil variation, and associated variation in learning, occurs in *H. m. malleti* and *H. m. rosina*. Future research should examine the neural responses of *H. m. malleti* and *H. m. rosina* males to an early social experience, as well as their relative flower color and nectaring location learning abilities and neural anatomy, to determine the mechanisms underlying their observed difference in response to prior social experience.

## Conclusion

Here we show that male *H. melpomene* butterflies use past social experience to inform current mating behavior. This response is lineage (wing pattern) specific, and coincides with lineage-specific differences in male assortative preference. Our findings strongly suggest that there are lineage-specific selective forces acting on cognitive function in *Heliconius* butterflies. Future research should explore the effect of cognition on speciation in this speciose group.

## Acknowledgements

We would like to thank Alexis Okoro for assistance with data processing, and Sushant Potdar, Grace Hirzel, Matt Murphy, Dylan Meyer, and Tim Sullivan for assistance with butterfly husbandry. This research was funded by the University of Arkansas.

## Data Accessibility Statement

Analyses reported in this article can be reproduced using the data provided by Rather et al., (XXX).

